# Cross-species analysis reveals unique and shared roles of Sox9 and Sox10 (SOXE family) transcription factors in melanoma

**DOI:** 10.1101/2022.12.05.519210

**Authors:** Eva T. Kramer, Paula M. Godoy, Charles K. Kaufman

## Abstract

SOX9 and SOX10 are two highly similar transcription factors with nearly 100% identity at their DNA binding domains. Both transcription factors play key but distinct roles in neural crest cell fate specification and melanoma formation. High expression of SOX9 and SOX10 appear to be mutually exclusive, with high SOX10 characteristic of proliferative melanoma and high SOX9 characteristic of metastatic melanoma. To further elucidate the role of SOX9 in melanoma, we over-express SOX9 in a zebrafish melanoma model and a human melanoma cell line. Analysis of tumor onset, binding dynamics, and transcriptional identities supports the notion of SOX9 driving a more mesenchymal signature, which is important for metastasis. Additionally, we identified a potential mechanism of SOX9 down-regulation via analysis of a functional and recurrent non-coding variant in human melanoma. Altogether, our results present a dosage-dependent role of SOX9 and, likely, SOX10 in the melanoma lifespan.

## INTRODUCTION

A persistent challenge in cancer biology arises when attempting to assign specific functions to genes with overlapping or potentially redundant functions. Cancers are notoriously adept at coopting normal cellular machinery and altering gene expression programs to produce more fit subclones to grow, to spread, and to evade innate and treatment-associated limitations on their survival. Melanoma has typified this highly plastic behavior, and efforts to understand the molecular details of such adaptations are made even more difficult when highly related genes and interdependent pathways are studied.

Cutaneous melanoma cancer typically arises from melanocytes within a highly UV-mutagenized field of melanocytes^1,2^. Reemergence of aspects of an embryonic neural crest program is a hallmark of melanoma initiation (White et al. 2011) with SOX10, a member of the SOXE family, highly expressed and functioning as a vital regulator of the reemergence of the neural crest in melanoma^3–6^. However, its closely related SOXE family member SOX9 is frequently downregulated in proliferative melanoma and, in some contexts, has an antagonistic effect on SOX10 expression when upregulated^7–9^.

SOX9 and SOX10 are members of the SOXE family and have similar structures, containing canonical high-mobility group (HMG), transactivation, homodimerization, and C-terminal domains with high degrees of sequence identity or similarity^10^. The highly conserved HMG domain has 79 amino acids which recognize CATTGT-like sequences^11^. Furthermore, SOX9 and SOX10 have similar DNA binding motifs, and thus can bind to the same DNA regions, yet regulate each region differently. SOX9 and SOX10 may also bind as monomers or dimers in different contexts, and they can heterodimerize at palindromic binding sites^10–13^.

SOX10 is expressed in migratory neural crest during the specification of the NC into melanocyte or glial lineages^14^. In zebrafish, the *sox9b* paralogue is expressed earlier than *sox10* in premigratory neural crest^7,14–17^. SOX9 is most associated as the master transcription factor for the chondrogenic lineage^18^. Nevertheless, overexpression of Sox9 in chick results in ectopic neural crest, with more differentiation towards melanocytes and glia (typically associated with *sox10*) and reduced differentiation away from the neuronal lineage and central nervous system^14^. Additionally, hypomorphic *sox9b* in zebrafish results in dispersed melanocytes and reduced iridophores, indicating it plays a role in melanocyte development as well^17^.

We and others have found that melanoma cells arising in the most widely used zebrafish melanoma model (BRAF/p53 dependent) show an upregulation of *sox10* and a downregulation of *sox9b* (the closest homolog to human SOX9) compared to melanocytes^19,20^. This raises the question of how these closely related transcription factors might function to regulate different downstream effects in melanocytes and melanoma, particularly in the process of melanoma initiation.

## RESULTS

### SOX9 is downregulated and local chromatin becomes less accessible in melanoma compared to melanocytes

To confirm alterations in key SoxE family transcription factors, SOX9 and SOX10, we compared gene expression of *sox9b* and *sox10* in zebrafish melanocytes versus melanoma and *SOX9* and *SOX10* in human nevi versus primary melanomas ^19,21^. Both cohorts show a decrease of sox9b/SOX9 expression in melanoma samples and a strong (in zebrafish) or a trend (in nevi versus melanoma) towards increase in sox10/SOX10 as expected (Figure 1A-1C). SOX9 does not have frequent point mutations in the coding region or associated structural variation in three of the largest sequencing melanoma cohorts: 5% in TCGA-SKCM cohort^22^, <1% in ICGC-MELA^23^, and 2.9% in the MSK IMPACT cohort^24^ (Figure 1D). Therefore, we looked for evidence of epigenetic changes at the *SOX9/sox9b* locus. Using previously published ATAC-seq obtained from *BRAF*^*V600E*^*/p53*^*-/-*^ zebrafish melanocytes and melanomas^19^, we observe a general decrease in accessible DNA surrounding the *sox9b* locus (Figure 1E).

**Figure 1.**
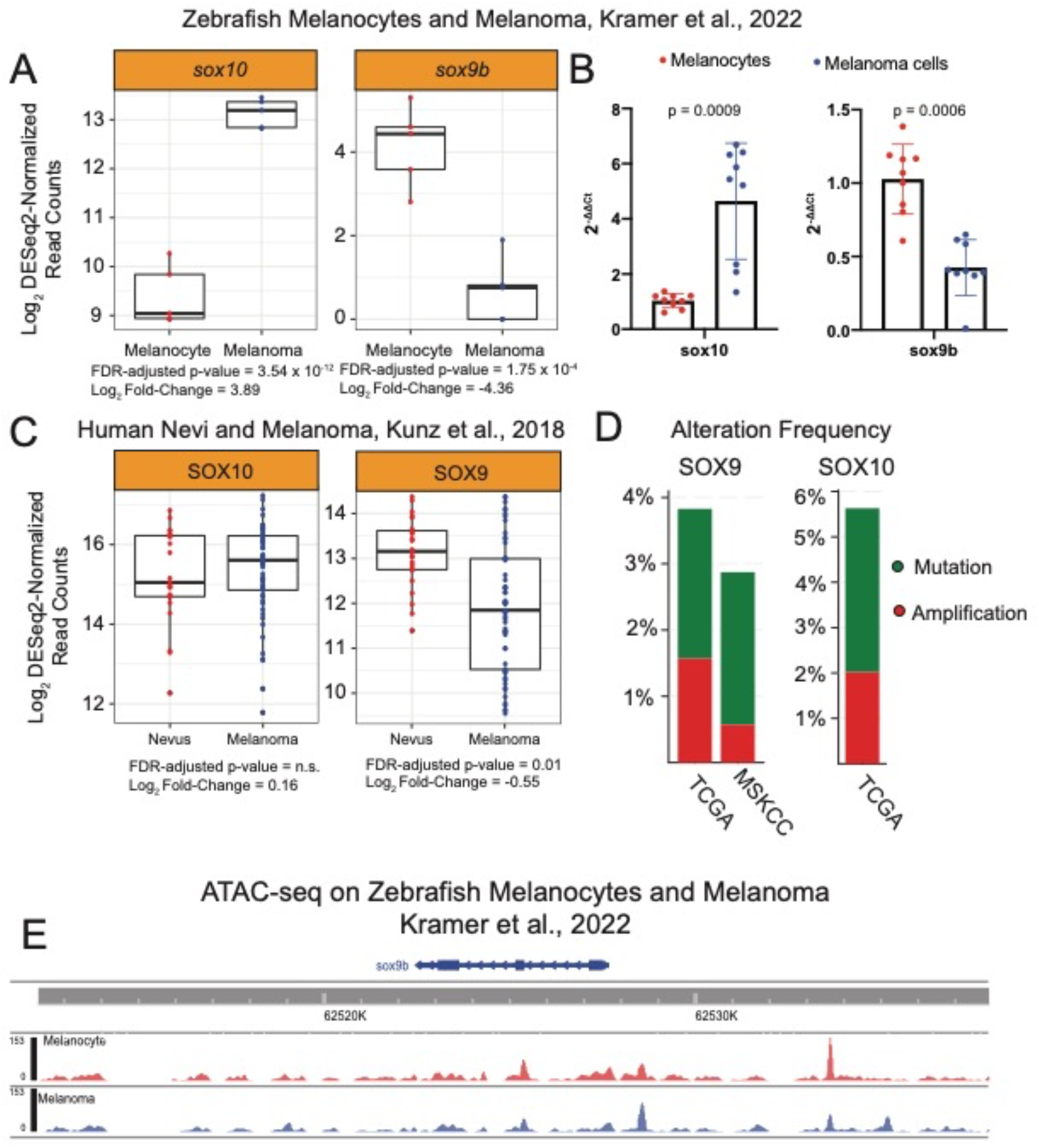
SOX9 and SOX10 alterations in melanoma. **(A)** Normalized read counts from RNA-sequencing and **(B)** qPCR for *sox10* and *sox9b* in melanocytes and melanoma cells from zebrafish with mCherry+ melanocytes and mCh+eGFP+ melanoma cells. Melanoma cells have an upregulation of sox10 and a downregulation of sox9b as measured by RNA-sequencing and qPCR. **(C)** Normalized read counts from a previously published RNA-sequencing dataset of primary melanomas and nevi. As in the zebrafish melanoma model, *SOX9* was up-regulated in nevi compared to primary melanomas. In contrast to the zebrafish melanoma model, *SOX10* was not significantly altered in nevi but trended higher in primary melanoma. **(D)** Alteration frequencies of SOX9 and SOX10 from the TCGA Pan Cancer Atlas and of SOX9 in the MSKCC IMPACT cohort. SOX10 was not assayed in the MSKCC cohort. **(E)** Genome tracks from the Epigenome Brower using combined bigwig files from ATAC-sequencing of mCh+eGFP+ melanoma cells from 8 tumors and mCh+ melanocytes from 4 zebrafish.

We then applied our recently published method to look for recurrently mutated non-coding regions in human melanoma and identified a mutation in the intron of *SOX9-AS1 (*chr17:70114939.70114939.G.A) that is detected in 6 out of 140 whole-genome sequenced cutaneous melanomas (4%, Figure 2A)^25^. We cloned a 300 bp region containing either the WT sequence or the recurrently mutated sequence upstream of a minimal promoter driving luciferase and observed a statistically significant decrease in reporter activity across all 3 melanoma cell lines assayed (Figure 2B). Motif analysis via motifBreakR^26^ predicts a motif gain of a GATA binding site (Figure 2C). Overall, we confirmed the down-regulation of *SOX9* and its zebrafish ortholog *sox9b* in two cohorts that specifically look at primary melanomas and nevi/melanocytes (Figure 1A-1C), identified both a decrease in accessibility in melanoma compared to melanocytes (Figure 1E), and a recurrent non-coding somatic variant in human melanoma that decreased reporter activity by putatively creating a binding site that represses transcription (Figure 2).

**Figure 2.**
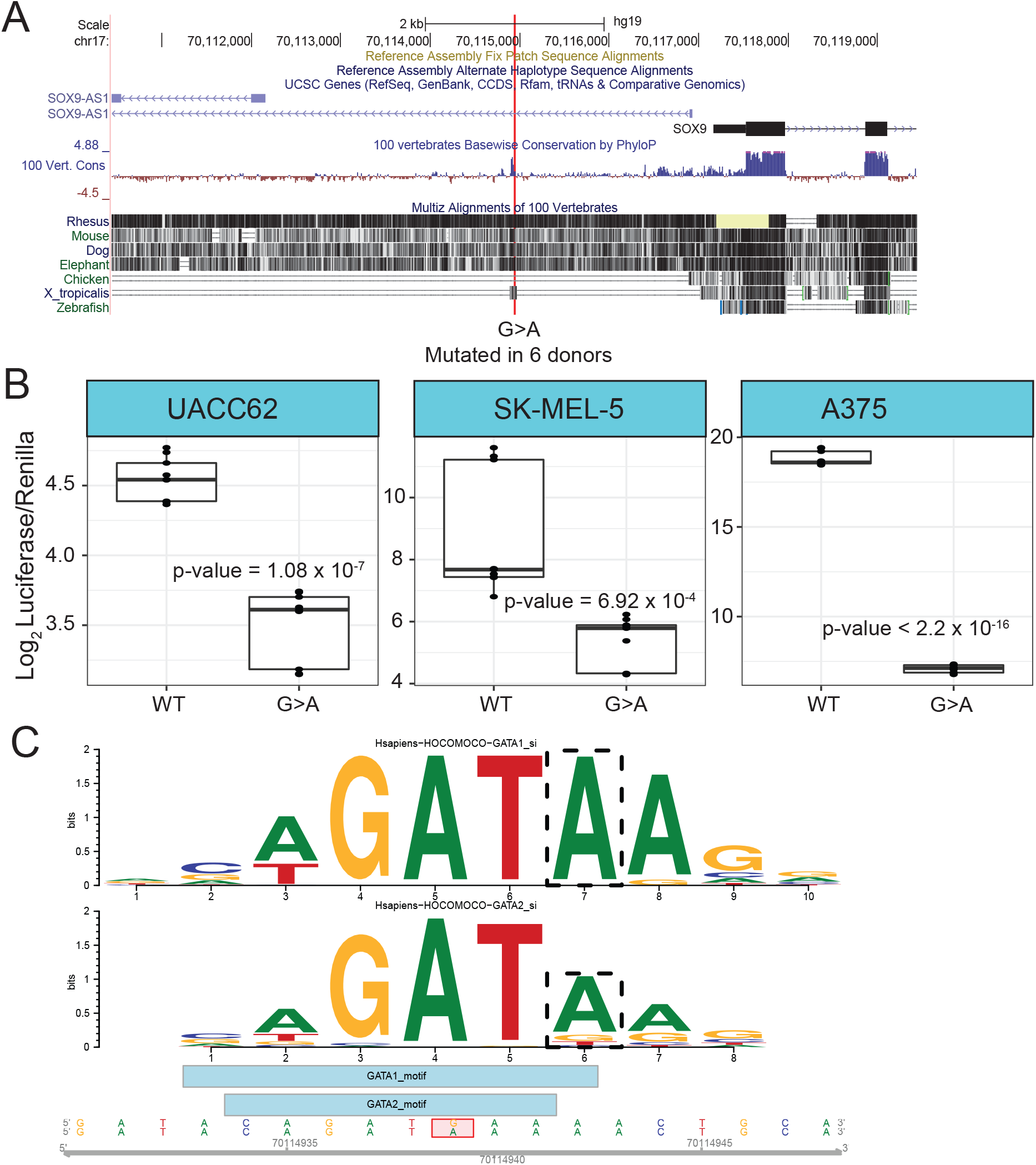
A recurrent non-coding variant in a putative SOX9 enhancer reduces luciferase activity. **(A)** Screenshot of UCSC Genome Browser capturing the SOX9 non-coding variant highlighted in red. A variant occurs within an intron of one isoform of the SOX9-AS1 gene, is approximately 2 kb from the transcriptional start site of SOX9, and is conserved across mammals and *X. tropicalis*. **(B)** Results from luciferase assay across three melanoma cell lines: A375, SK-MEL-5, and UACC62. Each point represents a log_2_-transformed luciferease value normalized to renilla. WT SOX9 enhancer sequences had higher activity than mutated (G>A) sequences. **(C)** A G>A variant in a putative SOX9 enhancer creates a novel GATA1/2 binding site. Position weight matrices of the GATA1 and GATA2 motif generated by the R package MotifBreakR. The variant is denoted by the dashed box.

### Overexpression of *sox9b* in melanocytes significantly slows melanoma onset

Based on the observation that SOX9/sox9b is decreased in melanomas relative to melanocytes, the cells-of-origin of melanoma, we asked whether over-expression of *sox9b* in a zebrafish melanoma model would slow melanoma onset. We used the widely applied miniCoopR system where zebrafish lines expressing the human BRAF^V600E^ oncogene in melanocytes (under control of the *mitfa* promoter) in a p53 null background are induced to develop melanomas by rescuing melanocyte formation with a plasmid carrying a mitfa-minigene and any gene of interest (also expressed under the control of the mitfa promoter)^27^. The miniCoopR plasmid (Figure 3A), which in this case contained either the *sox9b* or control *mCherry* gene is injected into zebrafish embryos with the successful integration of the miniCoopR plasmid yielding the rescue of melanocytes by three days post-fertilization (dpf). Zebrafish with rescued melanocytes are then tracked for the development of raised melanoma over time, allowing for determination of the length of time to tumor formation or % Tumor Free Survival.

**Figure 3.**
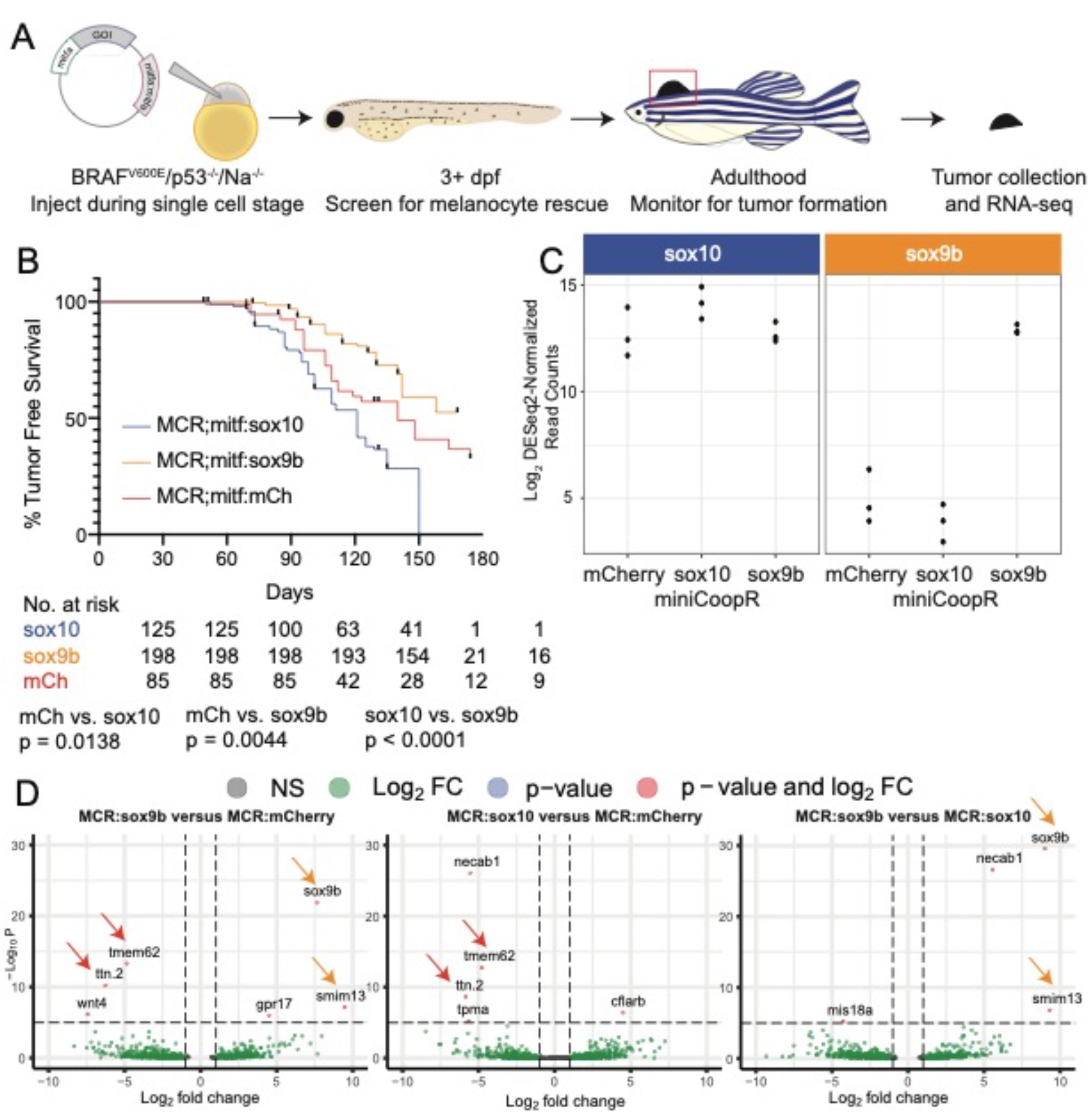
*sox9b* over-expression reduces melanoma onset in the miniCoopR zebrafish melanoma model. **(A)** Process for generating, injecting, and monitoring miniCoopR overexpression constructs in Tg(mitfa:BRAF^V600E^;p53^lf/lf^;Na^-/-^) zebrafish. **(B)** Overexpression of *sox9b* in a miniCoopR construct in a melanoma background results in slower melanoma onset (p = 0.0149) compared to *miniCoopR;mitfa:mCherry* and compared to *miniCoopR;mitfa:sox10* (p < 0.0001). Number of fish at risk for each condition listed underneath each date. **(C)** Log_2_-transformed DESeq2 normalized read counts of *sox10* and *sox9b* from RNA-sequencing of three tumors within each condition. *Sox9b* is upregulated only in the *miniCoopR;mitfa:sox9b* zebrafish whereas *sox10* has similar expression across all tumors. **(D)** Volcano plots depicting differentially expressed genes between all three conditions (FDR-adjusted p-value < 1 x 10^−5^ and Log_2_ Fold-Change > 1). Red arrows indicate genes consistently downregulated in sox9b/sox10 tumors compared to mCherry, and orange arrows indicate genes consistently upregulated in sox9b tumors compared to mCherry/sox10

Overexpression of *sox9b* significantly slowed melanoma onset (median onset 163 dpf) compared to *mCherry* (152 dpf, p = 0.0044) and *sox10* (125 dpf, p < 0.0001) while *sox10*, as previously reported^3^, sped melanoma onset (p = 0.0138) (Figure 3B). H&E staining of representative tumors show histologically grossly similar pigmented tumors in the tail of a miniCoopR:mCherry tumor (Supplemental Figure 1A) and in the dorsal area of a miniCoopR:sox9b tumor (Supplemental Figure 1B).

To identify changes in overall gene expression in melanomas with different transgenes, we performed RNA-sequencing of miniCoopR:sox9b tumors, miniCoopR:sox10 tumors, and control miniCoopR:mCherry tumors. These results show over-expression of *sox9b* in miniCoopR:sox9b tumors compared to miniCoopR:sox10 (log_2_ fold-change = 9.0, FDR-adjusted p-value = 1.67 x 10^−30^) and compared to miniCoopR:mCherry (log_2_ fold-change = 7.7, FDR-adjusted p-value= 1.36 x 10^−22^, Figure 3C, Supplemental Table 2). Interestingly, *sox10* was highly expressed across all conditions without significant increases in miniCoopR:sox10 in these *de novo* but established melanoma tumors, consistent with the dependency of melanoma on *sox10* in this zebrafish model (Figure 3C, Supplemental Table 2). Despite a large overexpression of *sox9b* expression in miniCoopR:sox9b compared to control miniCoopR:mCherry and miniCoopR:sox10 tumors, there were only 62 differentially expressed genes (FDR-adjusted p-value < 1 x 10^−5^ between miniCoopR:sox9b and miniCoopR:mCherry tumors, 54 genes between miniCoopR:sox10 and miniCoopR:mCherry tumors, and 57 genes between miniCoopR:sox9b and miniCoopR:sox10 tumors (only genes with FDR-adjusted p-value < 1 x 10^−5^ shown in Figure 3D, Supplemental Table 2).

Overall, our results suggest that over-expression of *sox9b* delays tumor onset, potentially through a gene regulatory network that is not captured by RNA-sequencing of bulk established tumors as performed here. While *sox9b* was only over-expressed in the miniCoopR:sox9b tumors, *sox10* is expressed at high levels across all fully formed tumors consistent with *sox10* up-regulation during tumorigenesis, as previously shown^3,6,19^.

### SOX9 over-expression in a human melanoma cell line displaces SOX10 but leads to no major alterations of the transcriptome

We next assayed the binding patterns of SOX9 and SOX10 in an established human melanoma cell line, A375, to begin to interrogate potential specific or shared roles of each factor. We over-expressed *SOX9* in A375, which has low endogenous transcript levels of *SOX9* and high *SOX10* compared to other melanoma cell lines and performed Cut&Run on H3K4me3 which mark actively transcribed promoters, SOX10, SOX10, SOX9 tagged by Myc, and SOX9 tagged by HA^28^. Although *SOX9* was over-expressed 12.2-fold (Figure 4A, FDR-adjusted p-value = 2.16 x 10^−204^), we only detected 86 differentially expressed genes (Supplemental Table 2). Of these few altered genes, however, CDH10, IGF2, and EHF belong to a gene signature associated specifically with a SOX9-high, mesenchymal-like state in melanomas^29^ (Figure 4B, in bold). In this context, *SOX9* overexpression did not suppress *SOX10* expression (Figure 4A), as in the established zebrafish tumors (Figure 3C). In line with a low number of differentially expressed genes, the distribution of H3K4me3 is similar in the A375 cell line over-expressing *SOX9* and in the WT A375 which has very low levels of *SOX9* (Figure 4C).

**Figure 4.**
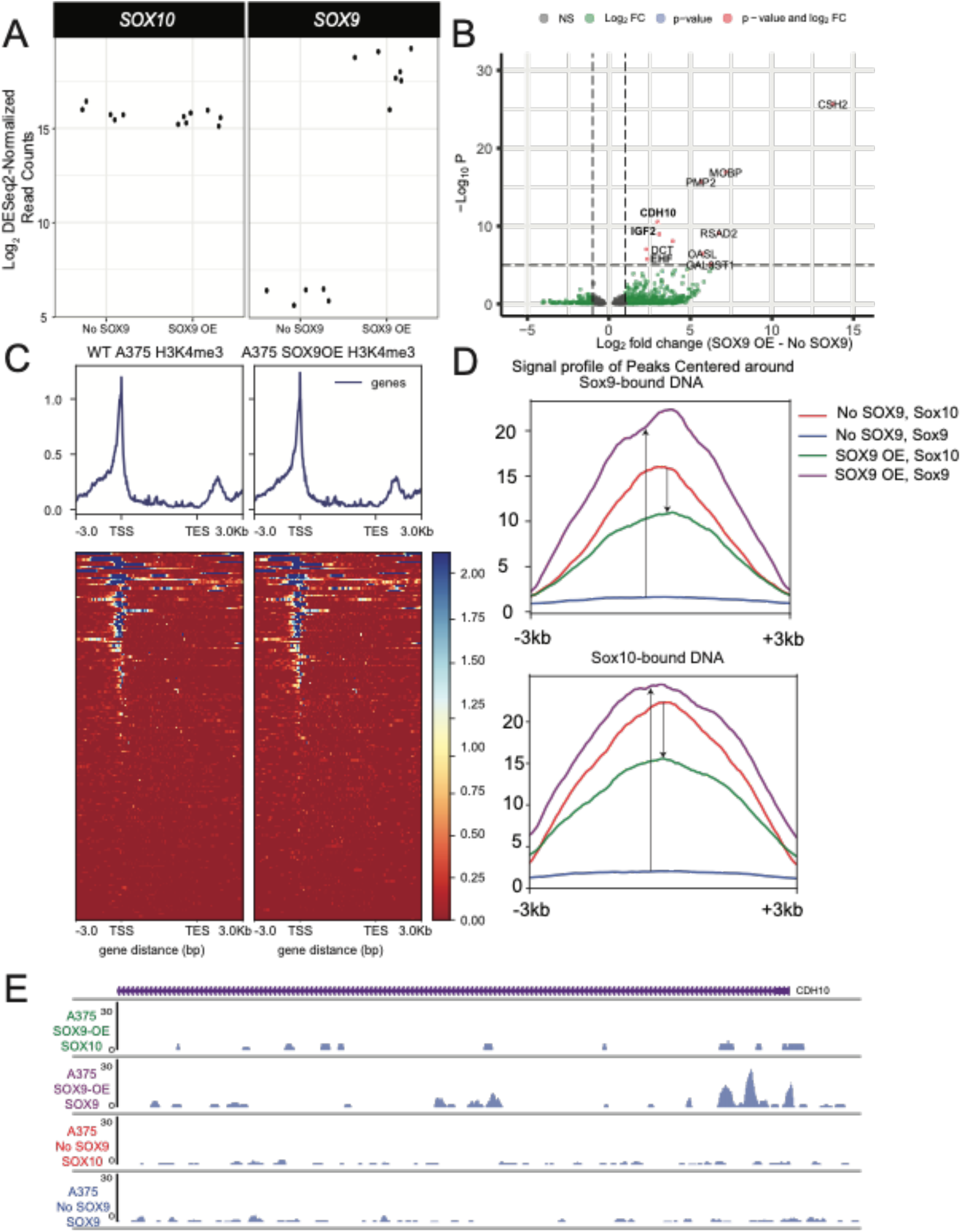
CUT&RUN analysis of SOX9 and SOX10 binding in SOX9 over-expressing A375 cells. **(A)** *SOX9* over-expression up-regulates *SOX9* in A375 but does not alter *SOX10*. Each point represents RNA-sequencing of either a WT A375 human melanoma cell line (No SOX9) or A375 with transient over-expression of *SOX9* (SOX9 OE). **(B)** Volcano plot depicting differentially expressed genes between No SOX9 and SOX9 OE conditions (log_2_ fold-change > 1 or < -1 and FDR-adjusted p-value < 0.05). **(C)** Density and heatmap profiles of H3K4me3 signal at the top 100 differentially expressed genes, ordered by FDR-adjusted p-value, in WT A375 or SOX9-OE A375. Each gene is scaled to the same length with an extra 3,000 base pairs upstream of the TSS and downstream of the TES. **(D)** Signal profiles of SOX9 and SOX10 binding at regions bound by SOX9 (top) and bound by SOX10 (bottom). Purple lines represent SOX9 binding in SOX9-OE cells, red lines represent SOX10 binding in WT cells, green lines represent SOX10 binding in SOX9-OE cells, and blue lines represent SOX9 binding in WT cells. Because WT A375 has endogenously low levels of SOX9, blue lines are near 0. Up arrows depict an increase in SOX9 binding between WT A375 and SOX9-OE A375. Down arrows depict a decrease in SOX10 binding upon SOX9-OE. **(E)** Screenshot of the Epigenome browser at the promoter of the mesenchymal-associated gene, CDH10. Bigwig files across each of the four conditions were combined across technical replicates.

We then asked whether the over-expression of *SOX9* led to SOX9 binding and whether this binding was cooperative with SOX10. Interestingly, we found that when *SOX9* was over-expressed (Figure 4D, blue line vs purple line), it would bind at native SOX10 binding sites (i.e. those locations where SOX10 binding was detected in the absence of SOX9 over-expression) displacing Sox10 (Figure 4D, red versus green line). Essentially, SOX10 did not as efficiently bind to DNA when *SOX9* was over-expressed. We observed similar binding sites via motif analysis (Supplemental Figure 2). In the case of the three mesenchymal-associated genes that were up-regulated upon *SOX9* over-expression, we noted only SOX9, but not SOX10, bound upstream at CDH10 (Figure 4E). Both SOX10 and SOX9 are bound to the EHF promoter but SOX9 appears to have higher levels of binding in this region (Supplemental Figure 3A). IGF2 did not have any detected binding activity at its promoter for either SOX9 or SOX10 (Supplemental Figure 3B).

Taken together, increased expression of *SOX9* did not drastically alter the gene expression program of established A375 melanoma cells, nor did it alter the distribution of H3K4me3 marks. However, SOX9 appears to bind at similar sites as SOX10, even displacing SOX10 in some cases, suggesting a potential for competitive interaction between SOX9 and SOX10, with potential specificity of SOX9 binding at some loci (e.g. near CDH10).

## DISCUSSION

Despite their high degree of similarity and potential interactions, SOX9 and SOX10 perform different roles in neural crest development and appear to have drastically different roles in melanoma emergence. Since SOX10 has been shown to promote melanoma formation in some contexts^3,5,6^, we investigated the more unclear role of SOX9 in melanoma. SOX9 has been implicated in damage response program as it is upregulated upon UV exposure and can activate key melanocyte genes^30^. However, upon melanoma onset in human and animal melanoma models, SOX9 becomes downregulated compared to melanocytes while SOX10 becomes upregulated^6,7,19,31^.

Although SOX9 is expressed at low levels in proliferative melanoma, it is enriched in metastatic melanomas, and overexpressing SOX9 leads to the activation of EMT pathways and an increase in the volume of metastatic tumors in mouse RAS melanoma models^8^. In human melanoma, *SOX9* is more associated with invasive signatures in contrast to *SOX10*, which is more expressed in proliferative melanoma signatures^32,33^. Dedifferentiated or mesenchymal subtypes of melanoma have higher SOX9 and lower SOX10 compared to more neural crest-like and differentiated melanomas^29,34^. Despite their structural similarities, SOX9 and SOX10 appear to contribute to melanoma formation via distinct mechanisms. In addition, a reasonable model supported by these and other published data^3,4,7^ continues to support that elevation of SOX10 activity supports early tumor formation/initiation, while SOX9-activity may support other aspects typically later functions of tumor behavior, e.g. metastasis. Our results also provide support for a less proliferative but more mesenchymal tumor subtype associated with *SOX9* overexpression^29,35^. Bulk analysis of RNA-sequencing of primary melanomas shows a general reduction of *sox9b/SOX9* in zebrafish and human melanomas. In support of this, overexpression of the zebrafish paralog *sox9b* significantly delayed tumor onset compared to overexpression of *mCherry* or *sox10*. Additionally, we identified a functional non-coding variant occurring in a SOX9 putative enhancer that reduces luciferase activity in three melanoma cell lines, suggesting a potential mechanism by which SOX9 is down-regulated in a subset of tumors, and could be speculated to support selection for a metastatic phenotype. Motif analysis of this variant shows the creation of a GATA1 site. Interestingly, GATA1 has been shown to repress the transcription of another SOX family member, *SOX2*, in a cancer stem cell line^36^.

As SOX9 and SOX10 have 96% similarity in their HMG domain which binds to DNA, we explored the binding dynamics and any associated transcriptional changes of SOX9 and SOX10 in A375, a human melanoma cell line, upon SOX9 over-expression^10^. Unexpectedly, we observed only twelve differentially expressed genes despite a 12-fold over-expression of *SOX9* and no associated changes in H3K4me3. Moreover, SOX9 appeared to bind precisely at SOX10 sites, displacing SOX10 in certain contexts. Although wild-type A375 has low levels of *SOX9*, the transcriptional identity of A375 is more mesenchymal than other melanoma cell lines, suggesting that SOX9 could act as a redundant transcription factor in this context. Interestingly, three of the twelve genes that were up-regulated in the *SOX9*-overexpressing cell lines were associated with high-expressing *SOX9* mesenchymal-like subpopulations from single-cell RNA-sequencing analysis of ten tumors^29^. Recently, single-cell RNA-sequencing of an NRAS-driven mouse melanoma tumor identified a small population of mesenchymal-like tumors that did not contribute to primary melanoma formation but were identified as metastasis-initiating cells^35^. These cells are suspected to be plastic, able to convert their transcriptional program to other identified clusters with gene regulatory networks that were more neural-crest like or undifferentiated-like. Transient changes in SOX9 and SOX10 may drive the plasticity between cluster subtypes, as knockdown of *SOX10* has been shown to lead to a more mesenchymal population that has higher migration capabilities^29^. Stabilization of either a SOX9 or SOX10 dominant cluster may require external input.

While further experiments are required to understand transient binding dynamics and associated transcriptional changes at different melanoma stages, our results are consistent with a dosage-dependent role for SOX9, where high levels of *SOX9* lead to a more mesenchymal subtype beneficial for initiating metastasis but not beneficial for driving primary tumor formation, during which low levels of *SOX9* are likely to be observed. These observations are important as therapies targeting SOX10 and SOX9 will need to consider potential compensatory mechanisms in metastatic melanomas.

## METHODS

### Analysis of SOX9 and SOX10 alterations

We downloaded normalized RNA-sequencing read counts from zebrafish melanocytes and melanoma cells and 23 laser-microdissected melanocytic nevi and 57 primary melanomas and plotted changes in expression with the R package *ggplot2*^19,21,37^. We used cBioPortal to look at mutation frequencies in the TCGA-SKCM and MSK-IMPACT cohorts^22,24,38^. Bigwig files corresponding to zebrafish melanocytes and melanoma cells were downloaded from GSE178803, uploaded onto the WashU Epigenome Browser^39^, and combined into one track representative of both conditions.

### qPCR of sox10 and sox9b in zebrafish

RNA was collected from cells using Trizol per manufacturer instructions. cDNA was prepared from total RNA using the SuperScript III First Strand Synthesis kit (ThermoFisher cat#18080044). Sybr Green Mastermix (Biorad cat#1725121) was used to perform qPCR in a Biorad CFX Connect Real-Time PCR System with triplicate technical replicates and 3 biological replicates. Primers are summarized in Supplemental Table 3.

### Melanoma Cell Culture

Melanoma cell lines were maintained at 37°C and 5% CO_2_ in 10 cm plates except where noted. A375 cells were obtained from and maintained as recommended by American Type Culture Collection (ATCC). A375 was cultured in DMEM + 10% fetal bovine serum (FBS). Media was supplemented with 100 mg/mL penicillin and 100 mg/mL streptomycin except during transfection.

### Validation of the SOX9 enhancer variant

Recurrent mutations upstream of SOX9 were identified using methods previously described^25^. For luciferase assays, we synthesized 300 bp sequences corresponding to WT and mutant hotspots with the variant centered at position 150 and ligated into a luciferase vector with a minimal promoter (pGL3-Promoter, E1761). Both the pGL3-Basic Luciferase vector (Promega, E1751) and the CDC20 promoter amplicon were digested using SacI-HF (NEB, R3156S) and XhoI (NEB, R0146S) at 37°C overnight, followed by heat inactivation at 65°C for 20 minutes. Digested vector and amplicon were ligated using T4 DNA Ligase (NEB, M0202S) and transformed into OneShot Top10 Chemically Competent Cells (ThermoFisher, C404010). Individual colonies were mini-prepped and confirmed by Sanger Sequencing (Azenta).

Using the Q5 Site-Directed Mutagenesis kit (NEB, E0554), we induced variants in the WT sequence using primers designed by NEBaseChanger (https://nebasechanger.neb.com/, Supplemental Table 3). Sequences that were successfully mutated, as well as the WT pGL3-Basic vector and pRL-TK (Promega, E2241), were midi-prepped (Qiagen, 12941).

For all transfections, 300,000 cells per well were seeded onto 6-well plates. All transfections were performed using 9 uL of Lipofectamine 2000 (Invitrogen, 11668), 1.5 μg of luciferase vector, and 1.0 μg of control pRL-TK (renilla), following the manufacturer’s protocol. All transfections were performed at minimum in duplicate. The following day, luciferase and renilla luminescence were measured using the Dual-Luciferase Reporter Assay System (Promega, E1910) per manufacturer specifications. Cells were lysed using 500 μL of 1X Passive Lysis Buffer and incubated for 15 minutes on an orbital shaker. 20 μL of lysate were added to clear-bottom 96-well plates. We ran three technical replicates per sample. Luminescence was measured on a GloMax 96 Microplate Luminometer (Promega) using a standard Dual Reporter Assay program. All luciferase values were normalized to renilla, as the internal transfection control. We then normalized all variant ratios to the corresponding average WT value. p-values were calculated using Student’s t-test.

### Zebrafish husbandry

Zebrafish were raised using standard animal protocols following Washington University IACUC. Embryos were generated with pairwise or harem crosses and raised at 28.5°C until 5-6dpf. Larvae were then transferred to a nursery where they were fed and raised with standard protocols associated with the Washington University Zebrafish Consortium. The following wild type, mutant, and transgenic strains were used: AB*, *Tg(mitfa:BRAF*^*V600E*^*); p53*^lf/lf^*;mitfa*^−/−^, *Tg(mitfa:BRAF*^*V600E*^*)/p53*^*lf/lf*^*/mitfa*^*+/-*^*/mitfa:mCherry, Tg(mitfa:BRAF*^*V600E*^*); p53*^lf/lf^*;crestin:eGFP, Tg(mitfa:BRAF*^*V600E*^*); p53*^lf/lf^*;crestin:eGFP;mitfa:mCherry*, and *crestin:eGFP*.

### Generating MiniCoopR overexpression constructs for candidate genes

The completed MCR plasmids were verified with restriction enzyme digestion and sequencing at GeneWiz to confirm the proper sequence was cloned. Previously created constructs were utilized for *sox10*^3^ and *mCherry*^19^. The construct for *sox9b* was cloned using Gateway cloning as previously described for all miniCoopR-based constructs^27,40^.

### Tumor onset curves

MiniCoopR constructs contain TOL2 sites that allow for integration of a gene of interest in the genome^41^. Each miniCoopR construct was injected into single cell Tg(*mitfa*:*BRAF*^*V600E*^); *p53*^-/-^; *mitfa*^-/-^ zebrafish embryos as previously described^27^. Larvae were screened for melanocyte rescue at 2-5 days post fertilization (dpf) to identify integration of the construct in the genome, then raised to adulthood and monitored for tumor onset. Starting at 6 weeks of age, rescued zebrafish were screened every 1-2 weeks to identify tumors. Data were entered into Prism and analyzed using Kaplan-Meier curves for tumor free survival with Mantel-Cox regression to determine p-values.

### RNA-sequencing of zebrafish tumors

RNA was collected from adult zebrafish tumors from fish overexpressing *miniCoopR;mitfa:sox9b, miniCoopR;mitfa:sox10*, and *miniCoopR;mitfa:mCherry* as previously described^19^. Sample preparation, sequencing, and analysis was performed by the Genome Technology Access Center using a Clontech SMARTer cDNA amplification kit. Sequencing was aligned to zv10 using STAR v2.0.4b^42^. Analysis was performed using the Genome Technology Access Center pipeline.

### Histology

Zebrafish were euthanized and fixed in 10% formalin and sent to HistoWiz for decalcification, embedding, and H&E staining of 5 sagittal sections which included the whole fish and/or the tumor.

### CUT&RUN

The following epitope tagged plasmids were obtained from GenScript using the Express Cloning service: SOX9-pcDNA3.1(+)-C-Myc and SOX9 OHu19789C_pcDNA3.1(+)-C-HA. 1 million cells were plated in a 10cm plate with normal media. The following day, complete media was replaced with media without antibiotics, and cells were transfected using 5 μg DNA + 15 μg PEI transfection reagent in Opti-MEM except where noted. After 48 hours, except where noted, cells were harvested for downstream processing.

CUT&RUN^28^ was performed using the Epicypher CUTANA ChIC/CUT&RUN kit (#14-1048) as written. Briefly, 500,000 cells were collected from each condition and bound to ConA beads. The cells were then incubated overnight at 4°C with 0.5 μg antibody (Supplemental Table 3) in an antibody buffer consisting of a cell permeabilization buffer composed of the Epicypher CUTANA pre-wash buffer with Roche cOmplete Mini EDTA-free protease inhibitor (#11836170001), 0.5mM spermidine (W1 additive in CUTANA v1.0 kit), and 0.01% digitonin (CP2 additive in CUTANA v1.0 kit), combined with 2 mM EDTA (AB3 Additive in CUTANA v1.0 kit). The following day, pAG-MNase was bound to the DNA, followed by chromatin digestion. DNA was purified and quantified on a Life Technologies Qubit 3.0 with a High Sensitivity Assay (Invitrogen, #Q32851).

Libraries were prepared with the NEBNext Ultra II DNA Library Prep Kit for Illumina (NEB, #E7645S) with NEBNext Multiplex Oligos for Illumina Dual Index Primers Set 1 (NEB, #E7600S) with primer indexing PCR run according to the Epicypher protocol. Libraries were assessed on an Agilent 2200 TapeStation with High Sensitivity D5000 ScreenTape and with a Qubit. Libraries were pooled together for each biological replicate (i.e., each round of transfection all samples were pooled together) and sequenced by the Washington University in St. Louis (WUSTL) Genome Technology Access Center (GTAC) with 50 million reads for the pool on a NovaSeq S4 System with 2x150bp read length.

GTAC generated demultiplexed fastq files for each sample. The quality of each file was assessed with FastQC. CutAdapt^43^ removed adapter sequences. Trimmed fastq files were aligned to hg38 using bowtie2^44^, then sorted with samtools^45^ and duplicates were removed with Picard. Peaks were called and normalized to IgG with MACS2^46^ with the following parameters: -f BAM -g hs –nomodel –pvalue 0.001. Output bam files were indexed with samtools. Then, bigwigs were generated using deepTools^47^ with the bamCoverage command and the following parameters: --extendReads 200 – ignoreDuplicates –minMappingQuality 10 –binSize 25 –scaleFactor 10 –normalizeUsing CPM.

Deeptools was used to generate profiles of H3K4me3 marks using a BED file of the gene start and gene end coordinates of the top 100 most differentially expressed ordered by FDR-adjuted p-values. To generate profiles of SOX9 and SOX10 binding, we used as input a BED file containing the location of the top 1000 peaks compared to IgG control using -log_10_p-value to order. Genome tracks were generated using the Epigenome Browser^39^.

### RNA-sequencing of cell lines

Total RNA was isolated from up to 1 million cells using Trizol (Thermo Fisher, #15596026) and quantified concentration with a Life Technologies Qubit 3.0 with a High Sensitivity Assay (Invitrogen, #Q32851). RNA was diluted to 10nM if the sample exceeded this concentration, then submitted to GTAC for quality assessment and quantification on an TapeStation 4200. RNA with RIN > 8.0 was subject to polyA selection (mRNA Direct kit, Life Technologies), fragmented, and cDNA was prepared with a SuperScript III reverse transcriptase enzyme. Adapters were ligated to the ends of ds-cDNA, then the library was prepared using unique dual index primers. The libraries were sequenced on an Illumina NovaSeq 6000 with 2x150bp read length.

Initial RNA-seq analysis was performed by GTAC. GTAC used Illumina’s bcl2fastq software and a custom demultiplexing program to perform basecalls and demultiplexing. Fastq files were aligned to hg38 (Ensemble release 76) using STAR. Gene counts were obtained using Subread featureCounts^48^. Total number of reads, number of uniquely aligned reads, and features detected were determined to evaluate sequencing quality. Gene counts were then imported into R. Normalization and differential gene analysis between conditions was performed using DESeq2^49^. Volcano plots were made using the R package, *EnhancedVolcano*.

## DATA ACCESS

RNA-sequencing and ATAC-sequencing of zebrafish tumors are available on the Gene Expression Omnibus (GEO) under accession number GSE178803. RNA sequencing of primary melanoma and nevi are available under accession number GSE112509. CUT&RUN and RNA-sequencing data is available under accession number GSE205046.

## COMPETING INTEREST STATEMENT

The authors declare no competing interest.

## ACKNOWLEDGEMENTS

Research reported in this publication was supported in part by the National Cancer Institute of the National Institutes of Health (NIH) under award number R01CA240633 and Melanoma Research Alliance #566840. The content is solely the responsibility of the authors and does not necessarily represent the official views of the NIH. C.K.K. was funded by the Cancer Research Foundation Young Investigator Award, NIH R01CA240633, and Melanoma Research Alliance #566840. E.T.K. was supported by T32# GM007067 and NIH R01CA240633. P.M.G. was supported by the National Science Foundation Graduate Research Fellowship (DEG-1745038) . We thank the Genome Technology Access Center in the Department of Genetics at Washington University School of Medicine for help with genomic analysis. The Center is partially supported by NCI Cancer Center Support Grant #P30 CA91842 to the Siteman Cancer Center and by ICTS/CTSA Grant# UL1 TR000448 from the National Center for Research Resources (NCRR), a component of the National Institutes of Health (NIH), and NIH Roadmap for Medical Research. Thank you to Anna Zarov and Brennan Lord-Howe for assistance with zebrafish analysis. Thank you to Megan Glaeser for perfecting the Western blots and qPCR. This publication is solely the responsibility of the authors and does not necessarily represent the official view of NCRR or NIH.

## SUPPLEMENTAL FIGURE LEGENDS

**Supplemental Figure 1.**
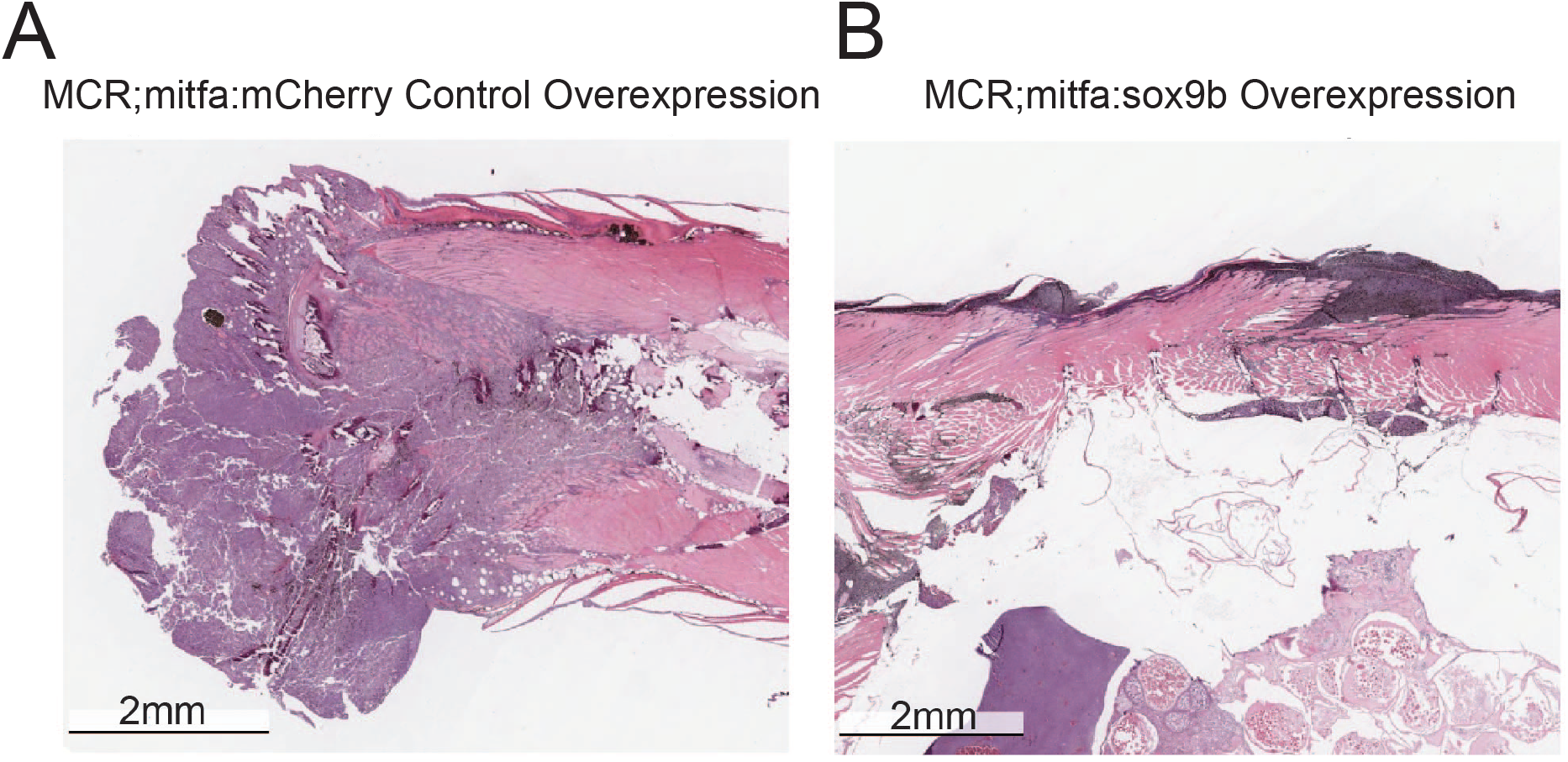
mCherry and sox9b tumors share similar histological features. Representative H&E histology images of Tg(BRAF^V600E^;p53^lf/lf^;mitfa^-/-^) zebrafish tumors injected with **(A)** miniCoopR;mitfa:mCherry and **(B)** miniCoopR;mitfa:sox9b. Tumor in **(A)** is in the tail while tumor in **(B)** is in dorsal area. Both tumors show similar invasion of the tumor into underlying muscle. MCR = miniCoopR.

**Supplemental Figure 2.**
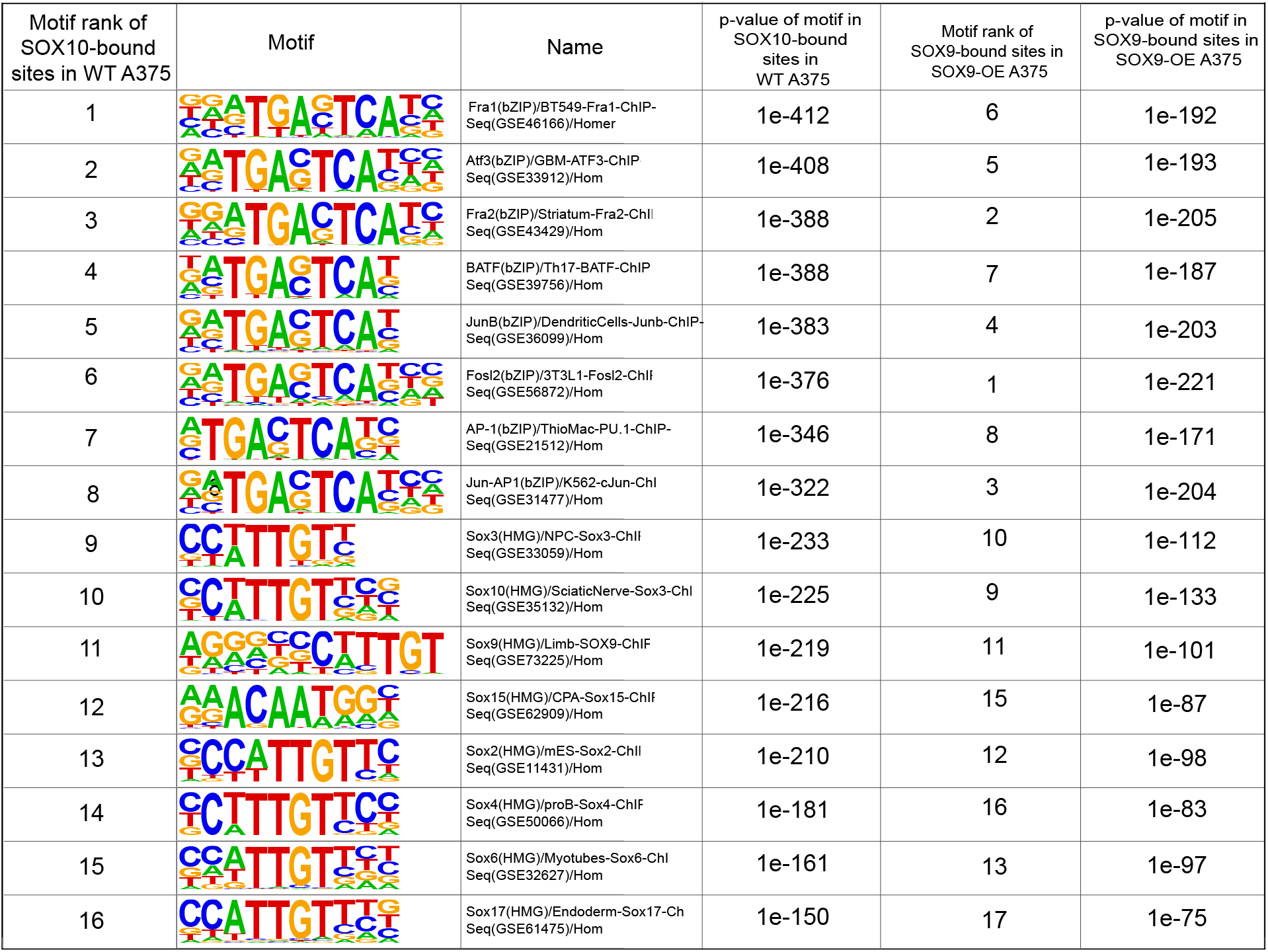
Motif enrichment at DNA bound by SOX9 and SOX10 in A375. Output from de novo motif analysis from HOMER. First column is the rank of the motif in SOX10 bound DNA in WT A375 (low endogenous levels of *SOX9*). The second and third columns provide details about the motif. The fourth column is the p-value associated with running motif enrichment of SOX10 bound DNA in WT A375. The fifth and sixth columns are the respective rank and p-values of motifs enriched in SOX9 bound DNA in A375 cells with transient *SOX9* over-expression.

**Supplemental Figure 3.**
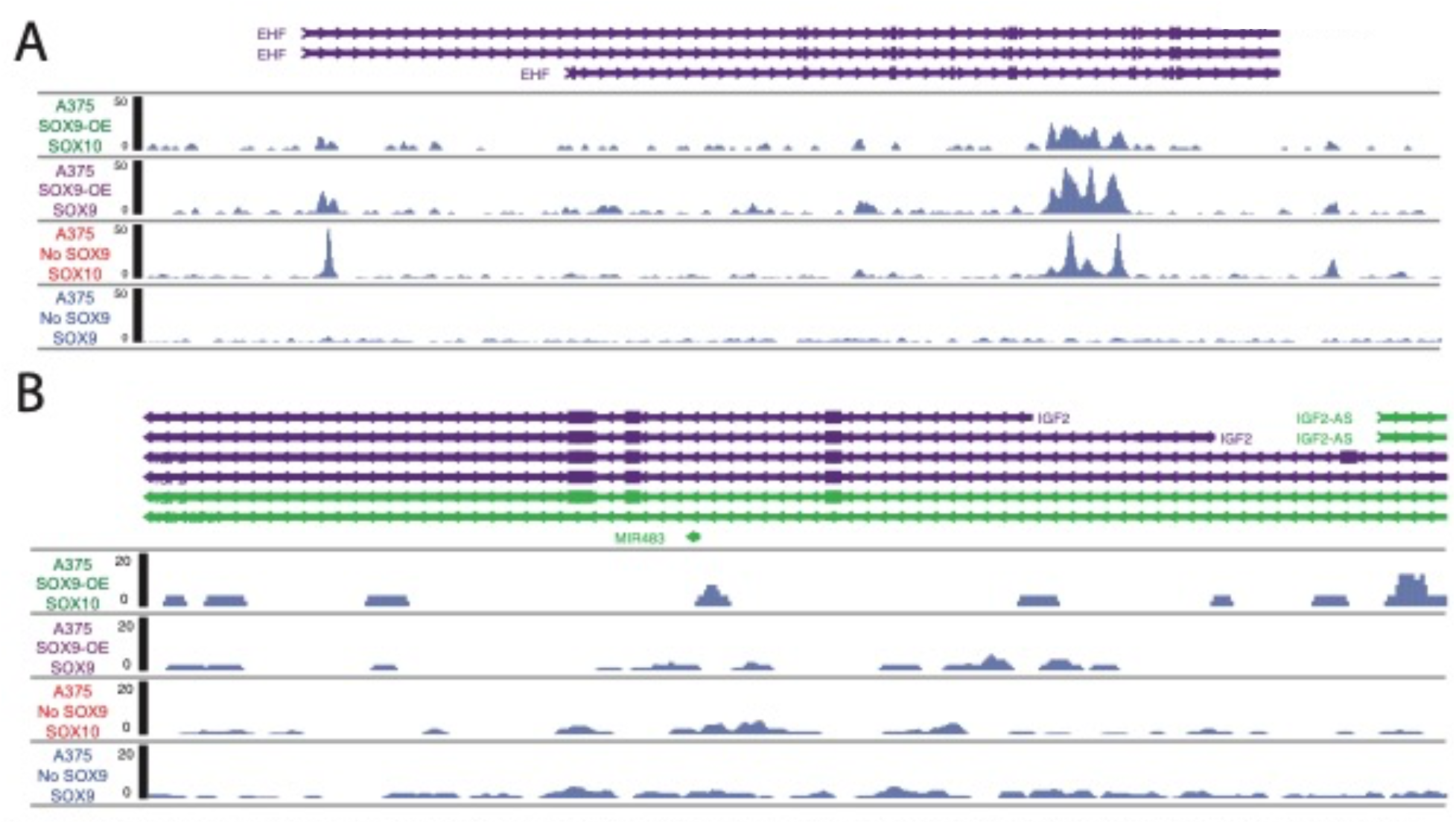
SOX9 and SOX10 binding at mesenchymal-associated genes. Screenshot of the Epigenome browser at the locus of the mesenchymal-associated gene **(A)** EHF and **(B)** IGF2. Bigwig files across each of the four conditions were combined across technical replicates.

## SUPPLEMENTAL TABLES

**Supplemental Table 1**. DESeq2-normalized read counts and results from differential expression analysis from miniCoopR:sox9b, miniCoopR:sox10, and miniCoopR:mCherry tumors.

**Supplemental Table 2**. DESeq2-normalized read counts and results from differential expression analysis from WT A375 (No SOX9) and SOX9-OE A375.

**Supplemental Table 3**. List of primers and antibodies used in this publication.

## Notes

### Competing Interest Statement

The authors have declared no competing interest.

