## Supplemental Figure 1 for "Cross-species analysis reveals unique and shared roles of Sox9 and Sox10 (SOXE family) transcription factors in melanoma"

A

MCR;mitfa:mCherry Control Overexpression

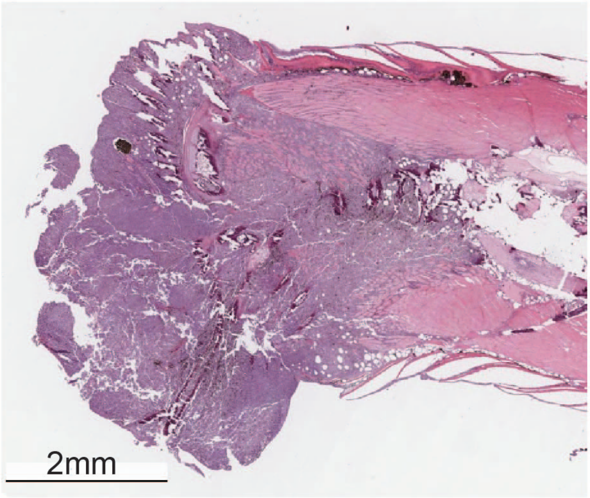


B

MCR;mitfa:sox9b Overexpression

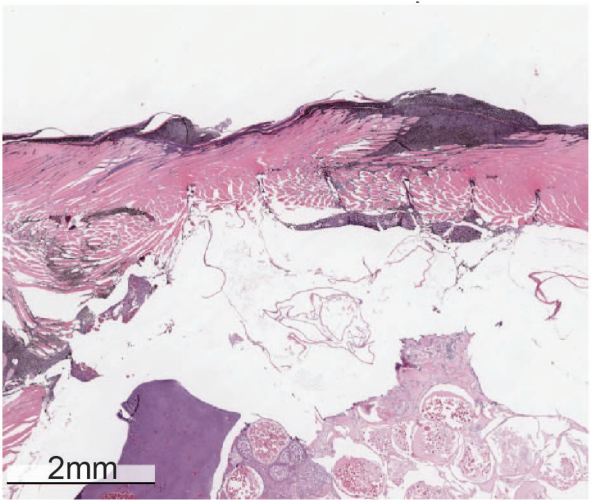
