## Supplemental Figure 2 for "Cross-species analysis reveals unique and shared roles of Sox9 and Sox10 (SOXE family) transcription factors in melanoma"

| Motif rank of SOX10-bound sites in WT A375 | Motif | Name | p-value of motif in SOX10-bound sites in WT A375 | Motif rank of SOX9-bound sites in SOX9-OE A375 | p-value of motif in SOX9-bound sites in SOX9-OE A375 |
|---|---|---|---|---|---|
| 1 | 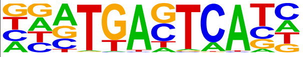 | Fra1(bZIP)/BT549-Fra1-ChIP-Seq(GSE46166)/Homer | 1e-412 | 6 | 1e-192 |
| 2 | 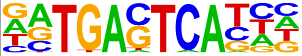 | Atf3(bZIP)/GBM-ATF3-ChIP-Seq(GSE33912)/Hom | 1e-408 | 5 | 1e-193 |
| 3 | 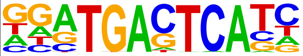 | Fra2(bZIP)/Striatum-Fra2-ChIl Seq(GSE43429)/Hom | 1e-388 | 2 | 1e-205 |
| 4 | 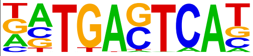 | BATF(bZIP)/Th17-BATF-ChIP Seq(GSE39756)/Hom | 1e-388 | 7 | 1e-187 |
| 5 | 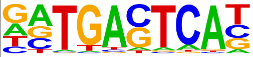 | JunB(bZIP)/DendriticCells-Junb-ChIP-Seq(GSE36099)/Hom | 1e-383 | 4 | 1e-203 |
| 6 | 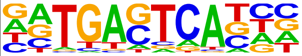 | Fosl2(bZIP)/3T3L1-Fosl2-ChIF Seq(GSE56872)/Hom | 1e-376 | 1 | 1e-221 |
| 7 | 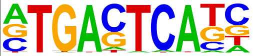 | AP-1(bZIP)/ThioMac-PU.1-ChIP-Seq(GSE21512)/Hom | 1e-346 | 8 | 1e-171 |
| 8 | 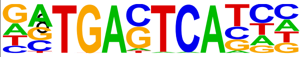 | Jun-AP1(bZIP)/K562-cJun-ChI Seq(GSE31477)/Hom | 1e-322 | 3 | 1e-204 |
| 9 | 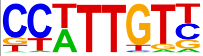 | Sox3(HMG)/NPC-Sox3-ChIF Seq(GSE33059)/Hom | 1e-233 | 10 | 1e-112 |
| 10 | 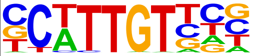 | Sox10(HMG)/SciaticNerve-Sox3-ChI Seq(GSE35132)/Hom | 1e-225 | 9 | 1e-133 |
| 11 | 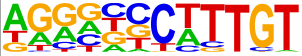 | Sox9(HMG)/Limb-SOX9-ChIF Seq(GSE73225)/Hom | 1e-219 | 11 | 1e-101 |
| 12 | 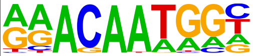 | Sox15(HMG)/CPA-Sox15-ChIF Seq(GSE62909)/Hom | 1e-216 | 15 | 1e-87 |
| 13 | 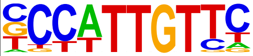 | Sox2(HMG)/mES-Sox2-ChIl Seq(GSE11431)/Hom | 1e-210 | 12 | 1e-98 |
| 14 | 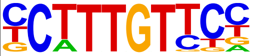 | Sox4(HMG)/proB-Sox4-ChIF Seq(GSE50066)/Hom | 1e-181 | 16 | 1e-83 |
| 15 | 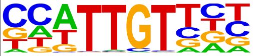 | Sox6(HMG)/Myotubes-Sox6-ChI Seq(GSE32627)/Hom | 1e-161 | 13 | 1e-97 |
| 16 | 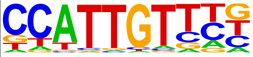 | Sox17(HMG)/Endoderm-Sox17-Ch Seq(GSE61475)/Hom | 1e-150 | 17 | 1e-75 |
