## Supplementary figures and images for "Cross-species analysis reveals unique and shared roles of Sox9 and Sox10 (SOXE family) transcription factors in melanoma"

### Supplemental Figure 3

Supplemental Figure 3

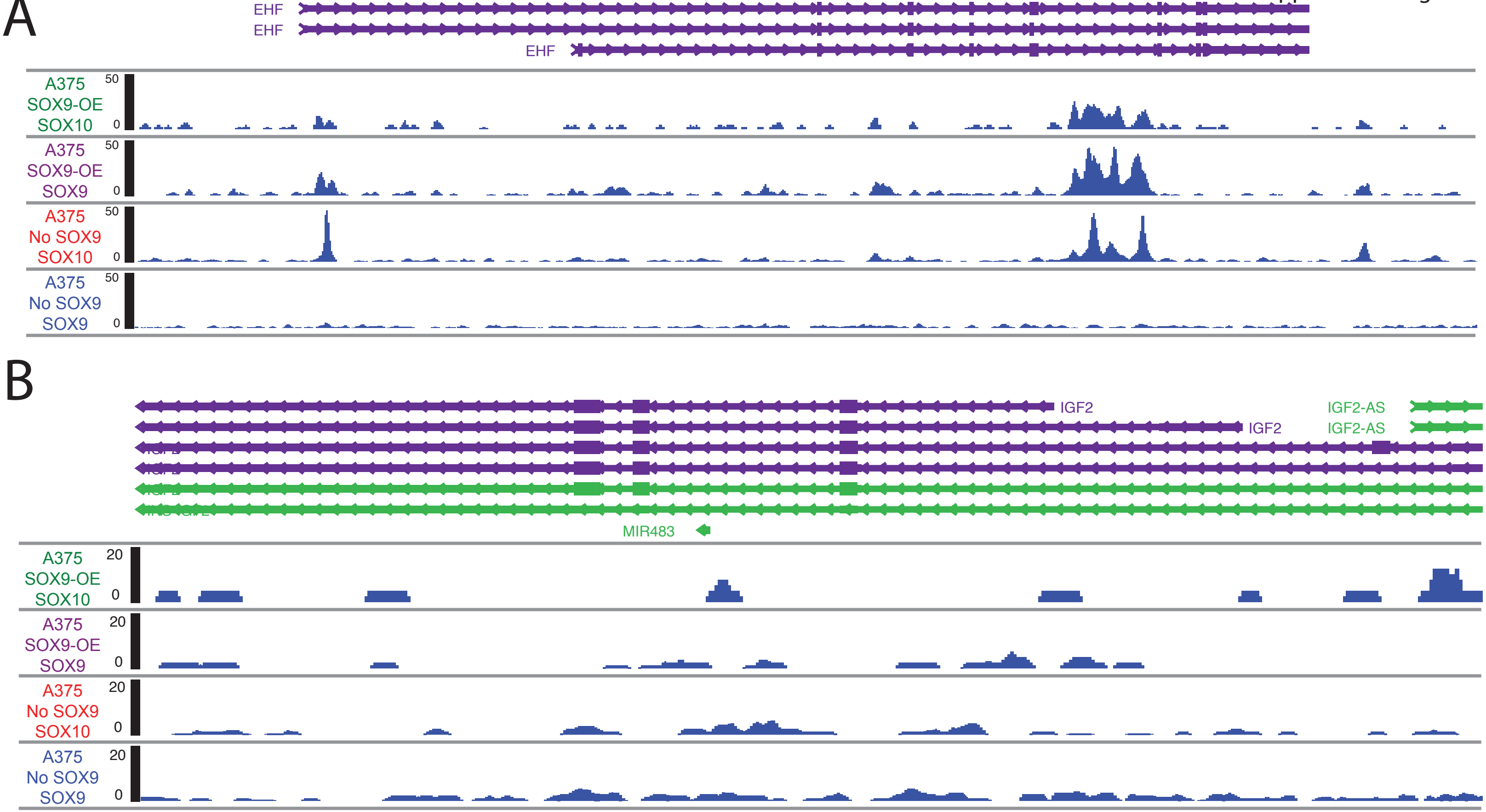
